# Design of nanoparticulate group 2 influenza hemagglutinin stem antigens that activate unmutated ancestor B cell receptors of broadly neutralizing antibody lineages

**DOI:** 10.1101/497123

**Authors:** Kizzmekia S. Corbett, Syed M. Moin, Hadi M. Yassine, Alberto Cagigi, Masaru Kanekiyo, Seyhan Boyoglu-Barnum, Sky I. Myers, Yaroslav Tsybovsky, Adam K. Wheatley, Chaim A. Schramm, Rebecca A. Gillespie, Wei Shi, Lingshu Wang, Yi Zhang, Sarah F. Andrews, M. Gordon Joyce, Michelle C. Crank, Daniel C. Douek, Adrian B. McDermott, John R. Mascola, Barney S. Graham, Jeffrey C. Boyington

## Abstract

Influenza vaccines targeting the highly-conserved stem of the hemagglutinin (HA) surface glycoprotein have the potential to protect against pandemic and drifted seasonal influenza viruses not covered by current vaccines. While HA stem-based immunogens derived from group 1 influenza A have been shown to induce intra-group heterosubtypic protection, HA stem-specific antibody lineages originating from group 2 may be more likely to possess broad cross-group reactivity. We report the structure-guided development of mammalian cell-expressed candidate vaccine immunogens based on influenza A group 2 H3 and H7 HA stem trimers displayed on self-assembling ferritin nanoparticles using an iterative, multipronged approach involving helix stabilization, loop optimization, disulfide bond addition, and side chain repacking. These immunogens were thermostable, formed uniform and symmetric nanoparticles, were recognized by cross-group-reactive broadly neutralizing antibodies (bNAbs) with nanomolar affinity, and elicited protective, homosubtypic antibodies in mice. Importantly, several immunogens were able to activate B cells expressing inferred unmutated common ancestor (UCA) versions of cross-group-reactive human bNAbs from two multi-donor classes, suggesting they could initiate elicitation of these bNAbs in humans.

**Importance:** Current influenza vaccines are primarily strain specific, requiring annual updates and offer minimal protection against drifted seasonal or pandemic strains. The highly conserved stem region of hemagglutinin (HA) of group 2 influenza A subtypes is a promising target for vaccine elicitation of broad cross-group protection against divergent strains. We used structure-guided protein engineering employing multiple protein stabilization methods simultaneously to develop group 2 HA stem-based candidate influenza A immunogens displayed as trimers on self-assembling nanoparticles. Characterization of antigenicity, thermostability, and particle formation confirmed structural integrity. Group 2 HA stem antigen designs were identified, that when displayed on ferritin nanoparticles activated B cells expressing inferred unmutated common ancestor (UCA) versions of human antibody lineages associated with cross-group reactive, broadly neutralizing antibodies (bNAbs). Immunization of mice led to protection against a lethal homosubtypic influenza challenge. These candidate vaccines are now being manufactured for clinical evaluation.

## INTRODUCTION

Influenza continues to be a significant global health burden, typically resulting in about 500,000 deaths worldwide annually (1), even though the technology for conventional egg-grown, whole inactivated influenza virus vaccines was developed more than 70 years ago. Constant antigenic drift of the influenza virus hemagglutinin (HA) coupled with immunodominant strain-specific antibody responses directed to the variable HA head domain results in conventional vaccine effectiveness ranging from 10-60% (2) and the need for seasonal updates of virus strains included in licensed vaccines. Furthermore, current vaccine approaches provide minimal protection against pandemic influenza strains (3, 4). Improved influenza vaccines would not be produced in eggs, would be designed to induce broad immunity against future drifted and pandemic strains without seasonal reformulation, and elicit durable immune responses avoiding the need for annual vaccination (4). One approach for achieving broadly cross-reactive influenza-specific antibodies is to target highly conserved neutralization-sensitive epitopes in the stem region of the influenza A HA surface glycoprotein (5). Monoclonal antibodies have been identified that bind conserved HA stem epitopes and possess broad neutralizing activity across diverse HA subtypes within an influenza A group, and some that have cross-group neutralizing activity (6–16). We and others have recently developed influenza A group 1 HA stem trimer antigens (17, 18). One candidate displaying H1-based HA stem trimers on self-assembling ferritin nanoparticles protected mice and ferrets from a lethal heterosubtypic H5N1 influenza challenge (17). However, these group 1 HA stem immunogens did not provide cross-group protection against group 2 viruses such as H3N2 and H7N9. The need for a better group 2 vaccine is particularly acute as vaccine effectiveness against H3N2 over the past decade has averaged only 33% (19), recent H3N2 strains have exhibited increased virulence (20), and H7 viruses represent one of the greatest pandemic threats from non-seasonal strains. H7N9 immunization was recently shown to induce multiple HA stem-directed antibody lineages in humans that recognized both group 1 and group 2 HA molecules, whereas H5N1 immunization induced primarily HV1-69 antibody lineages that cross-reacted primarily within group 1 influenza A subtypes (13). This suggests that group 2 HA stem immunogens may be more effective at inducing cross-group responses than group 1 immunogens in humans with pre-existing influenza immunity, or priming multi-donor antibody lineages with the capacity for broad neutralizing activity in influenza-naïve infants. In our studies, we used iterative structure-based design to develop headless group 2 HA stabilized stem trimers displayed on ferritin nanoparticles based on hemagglutinin sequences from H3N2 (H3ssF) and H7N9 (H7ssF) viruses as candidate vaccines. Notably, structural subtleties ultimately required the use of a completely different set of design strategies to stabilize group 2 HA stem immunogens relative to simpler hydrophobic repacking used to stabilize group 1 immunogens. Ultimately, H3ssF and H7ssF immunogen designs were identified that could activate B cells expressing inferred unmutated common ancestor (UCA) HA stem-directed human broadly neutralizing antibodies (bNAbs), suggesting the potential for inducing cross-reactive antibody lineages with the capacity to develop broad influenza neutralizing activity in humans.

## RESULTS

### Design of group 2 stabilized HA stem nanoparticle immunogens

Designing group 2 stabilized HA stem nanoparticles incorporated some features of group 1 H1-stabilized HA stem nanoparticle immunogens (17) (referred to as H1ssF herein). This comprised replacing the HA head region (residues HA1 43-313, H3 numbering (21)) with a GSG loop, replacing the HA2 interhelical region (residues 60-92) with a six-residue glycine-rich loop, including two repacking substitutions in the HA2 hydrophobic core (K103M/E51L), and connecting the C-terminal HA2 residue 174 to the N-terminal residue 5 of bacterial ferritin with a short linker (Fig. 1A and Figs. S1-2). In the group 1 context this strategy was sufficient for displaying antigenically correct H1, H2 and H5 HA stem trimers on ferritin nanoparticles (17). However, multiple additional steps were needed to achieve expression of group 2 H3 and H7 HA stem nanoparticles in mammalian HEK293 cells (Fig. 1). Although there is structural similarity between H1 and H3 HA stem regions (e.g. root mean square deviation (RMSD) of 1.0 Å for 134 stem residues between A/California/04/2009 (H1N1), PDB 3UBQ and A/Victoria/361/2011 (H3N2), PDB 4O5N) and sequence identity of approximately 55%, phylogenetic analysis indicates groups 1 and 2 comprise two distinct branches with consistent and predictable amino acid differences even for the stem region of HA (Fig. 1B). We hypothesized that further structural stabilization was required for expression and therefore, an iterative structure-based design effort was employed to stabilize the HA stem region of H3ssF immunogens using the 1.9 Å resolution crystal structure of HA for A/Finland/486/2004 (H3N2), (PDB 2YP2) (22) as a template. To screen for expression and correct antigenicity, designs were expressed in HEK 293T cells using a high-throughput 96-well format (23). Supernatants containing HA stem trimers genetically fused to ferritin nanoparticles were assessed directly for antigenic recognition by either an immunoprecipitation assay using the human cross-group, HA stem-specific bNAb FI6v3 or by an ELISA using the bNAbs CR9114, FI6v3, and CT149. CT149 was specifically included to assess trimer integrity because its epitope incorporates two adjacent protomers of the HA stem and thus may be sensitive to quaternary structure (15).

**Fig. 1.**
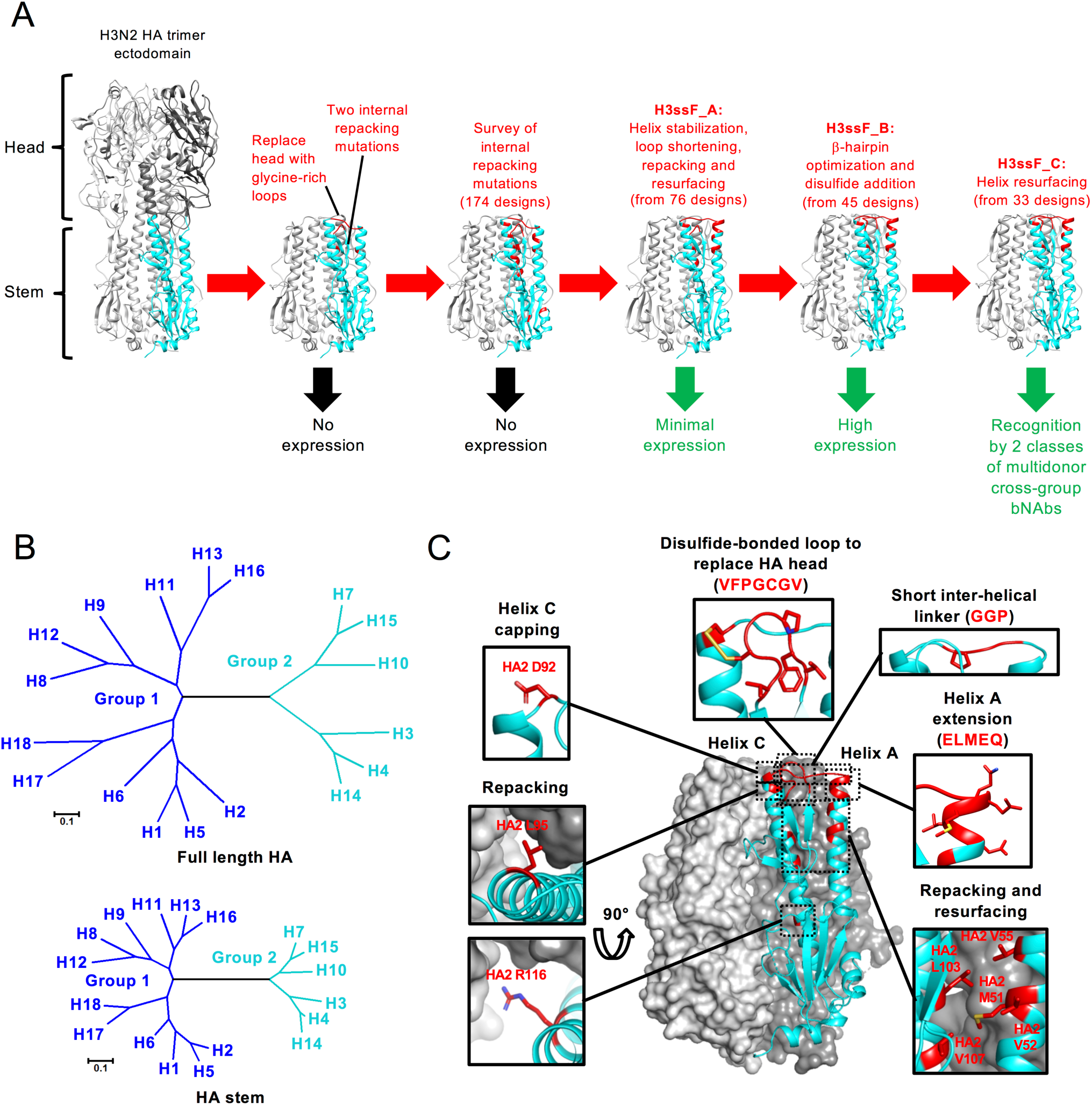
Structure-based design of group 2 HA stem nanoparticles. (A) Ribbon models depicting the HA stem nanoparticle design pathway for stabilizing the HA stem starting from an H3N2 HA trimer ectodomain. Two of the three protomers are gray and one protomer is cyan with the design modifications in red. Positive or negative outcomes are indicated by green or black arrows, respectively, below each step. bNAb, broadly neutralizing antibody. (B) Maximum likelihood phylogenetic tree for protein sequences from representative full-length HAs (top panel) and HA stem (bottom panel) of the 18 different subtypes. Group 1 subtypes are navy blue and group 2 subtypes are cyan. (C) A molecular model of the HA stem of an H3ssF_C design is shown colored as in (A). Two of the three protomers are depicted as surface representations and one protomer is displayed as a ribbon diagram. Insets show close-ups of the modified regions with stick representations for side chains of modified residues. Surface representations are removed from some insets for clarity. PDB entry 2YP2 was used an H3N2 model template.

Since internal hydrophobic repacking mutations were critical for the stabilization of the group 1 HA stem immunogens (17), and the burying of just a single -CH_2-_group in a protein interior can add approximately 1.1 kcal/mol of free energy to protein stability (24, 25), we explored additional repacking mutations selected computationally using Rosetta (26). In combination with some of the repacking mutations we also explored limited residue swapping between H1N1 and H3N2 subtypes (Table S1). Surprisingly, none of these 174 mutants could be expressed and recognized by bNAbs CR9114 or FI6v3 (Table S1), indicating novel approaches for structural stabilization would be required.

Upon further investigation of multiple protein stabilization strategies involving another 76 designs (Table S1), a lead design was successfully expressed (H3ssF_A), that was recognized by bNAb HA stem antibody FI6v3 by immunoprecipitation, and self-assembled into nanoparticles (Table 1, Table S1, and Figs. S2-3). H3ssF_A included the initial H1ssF-based mutations in addition to: i) helix stabilization, ii) loop optimization, and iii) point mutations of select H3 residues to improve internal packing (Figs. 1A and 1C and Fig. S2A). Each of the successful alterations were focused on the membrane distal end of the two central helices (A and C) of the HA2 fold. The outer helix A in H3N2 HA is approximately five residues shorter at its C-terminus than in H1 HA and it is known that helix length is related to stability (27). Therefore, helix A was extended at its C-terminus by five residues (ALMAQ) with high helix-forming propensities (28). Likewise, the N-terminus of the adjacent helix C was stabilized by adding an N-terminal Asp (HA2 92) as a helix-capping residue (29, 30) and two other mutations (HA2 S93A and N95L) at the N-terminus of helix C were made to increase its helix-forming propensity and improve interprotomer packing (Fig. 1C). Biophysical studies have shown that excess loop length can destabilize protein folds by up to several kcal/mol (31), and in the initial H1ssF and H3ssF designs, helices A and C were connected by a six-residue linker even though they are separated by only 12 Å. Therefore, the glycine-rich linker was shortened to a four residue GGPD linker to more closely match the interhelical distance. Lastly, four residues (HA2 52, 55, 107 and 116) on helices A and C respectively, were mutated to improve side chain packing (Fig. 1C and Fig. S2A). Three of these (HA2 52, 55 and 116) comprised mutations to H1 residues.

**Table 1.** Physical characteristics of the H3ssF and H7ssF nanoparticles

| Subtype | Design | Design characteristics |  |  |  |  | Relative Yield<sup>f</sup> | Tm (°C) | EM characterization |  |
| --- | --- | --- | --- | --- | --- | --- | --- | --- | --- | --- |
|  |  | HA1 loop<sup>a</sup> | Helix A extension<sup>b</sup> | Inter-helix loop<sup>c</sup> | Helix C mutations<sup>d</sup> | Predicted disulfides<sup>e</sup> |  |  | Ferritin core size (Å) | HA stem size (Å) |
| H3 | H3ssF_A | --GSG-- | ALMAQ | GGP | DAYL | NA | - | ND | ND | ND |
| H3 | H3ssF_B | VFPGCGV | ALMAQ | GGP | DCYL | DS1 | +++ | 66.2 | 124.0±7.0 | 82.0±9.0 |
| H7 | H7ssF_B | VFPGCGV | ALMAQ | GGP | DCYL | DS1 | +++ | 58.6 | 126.9±6.8 | 71.6±6.5 |
| H3 | H3ssF_C | VFPGCGV | ELMEQ | GGP | DCYL | DS1 | ++ | 66.6 | 123.2±3.4 | 65.0±5.2 |
| H7 | H7ssF_C | VFPGCGV | ELMEQ | GGP | DCYL | DS1 | +++ | 61.6 | 124.4±4.4 | 66.9±6.4 |
| H3 | H3ssF_D | VFPGC-V | ALMAQ | GGP | DCYL | DS1 | ++ | 66.3 | 125.3±5.1 | 67.7±12.4 |
| H7 | H7ssF_D | VFPGC-V | ALMAQ | GGP | DCYL | DS1 | +++ | 61.2 | 139.7±4.8 | 64.8±3.4 |
| H3 | H3ssF_E | VFP-CGV | ALMAQ | GGP | DCYL | DS1 | ++ | 63.0 | 130.1±5.3 | 66.9±7.2 |
| H7 | H7ssF_E | VFP-CGV | ALMAQ | GGP | DCYL | DS1 | + | 56.7 | 124.7±6.5 | 68.1±2.7 |
| H3 | H3ssF_F | VFPGCGV | ALMEE | GGP | DCYL | DS1 | + | 65.6 | 127.1±6.0 | 67.0±4.1 |
| H7 | H7ssF_F | VFPGCGV | ALMEE | GGP | DCYL | DS1 | +++ | 59.9 | 130.2±6.3 | 65.1±5.5 |
| H3 | H3ssF_G | CFNGIC- | ALMAQ | GGP | DAYL | DS2 | +++ | 59.9 | 126.3±4.0 | 68.5±3.3 |
| H3 | H3ssF_H | VFPGCGV | ELMEQ | GGP | DCYL | DS1, DS3 | ++ | 56.6 | 126.9±3.5 | 67.8±4.5 |
| H7 | H7ssF_I | VFPNCGV | ALMAQ | GGP | DCYL | DS1 | +++ | 58.7 | 124.3±4.1 | 68.4±6.6 |
| H7 | H7ssF_J | VFPGCGV | ALMAQ | GPP | DCYL | DS1 | +++ | 57.0 | 131.4±5.1 | 62.6±7.6 |

<sup>a</sup>Between HA1 residues 42 and 314 (H3 numbering)
<sup>b</sup>Directly following HA1 56
<sup>c</sup>Directly following the helix A extension
<sup>d</sup>Region between the interhelix loop and HA2 96
<sup>e</sup>DS1, HA1 C145-HA2 C93; DS2, HA1 C141-C146; DS3, HA1 C23-C322; NA, not applicable.
<sup>f</sup>Based on gel electrophoresis analysis of unpurified supernatants following transient transfection.

To further improve expression and stability, another 45 designs were explored, focusing on the short β-hairpin connected to a three-residue loop which replaced the HA head in HA1 (Fig. 1). Since the stability of short, two-stranded β-sheets has been observed to increase with length (32), we replaced the GSG loop with longer loops of 6-7 residues designed to double the hairpin length. Residues with high strand-forming propensities (28) such as Val, Ile and Phe were used for strand extension and dipeptides with propensities for form β-turns (33, 34) such as PG and NG were used to connect them. Disulfides were also designed into the hairpin loops for added stability (35). From an initial survey of loop extensions, H3ssF_B, designed as a 15-residue hairpin loop disulfide-bonded to the N-terminus of helix C resulted in expression levels of greater than 5 mg/L and recognition by CT149 (Fig. 1, Table 1 and Table S1).

Further loop optimization of the H3ssF_B design included removing a glycine from either side of the cysteine in the β-hairpin loop (H3ssF_D and H3ssF_E) and incorporating a different β-hairpin sequence with an internal disulfide bond engineered to link both strands together (H3ssF_G). Additional H3ssF_B variants incorporated hydrophilic, helix preferring Glu into the relatively hydrophobic helix A extension motif for improved solubility (H3ssF_C, H3ssF_F and H3ssF_H), and an engineered intra-protomer disulfide between HA1 residues 23 and 332 (H3ssF_H). All six variants resulted in nanoparticles recognized by CT149 in ELISA assays, but did not improve overall expression beyond that of H3ssF_B. Based on the successful expression and antigenic recognition of multiple H3ssF variants, we transferred mutations from constructs H3ssF_B, H3ssF_C, H3ssF_D, H3ssF_E, and H3ssF_F into analogous H7ssF nanoparticles using sequences from the human H7N9 strain (A/Shanghai/2/2013) and also created two additional loop variants (H7ssF_I and H7ssF_J) (Fig. S2B). All seven of these H7ssF designs expressed milligram per liter quantities of nanoparticles recognized by CT149 (Table 1).

### Physical characterization of H3ssF and H7ssF nanoparticles

Fourteen nanoparticles were selected for further characterization. These were purified by lectin chromatography followed by size exclusion chromatography, which resulted in distinct peaks consistent with approximately 1.2 MDa particles (Fig. 2A-B and Figs. S4). Negative stain electron microscopy of purified nanoparticles followed by reference-free 2D classification and averaging revealed spherical particles with five to six visible regularly-spaced protruding spikes (Figs. 2C-D, 2E-F and Figs. S4). All nanoparticles contained ferritin cores consistent with the expected 12-13 nm diameter, and HA stems with the expected length of approximately 7 nm (Table 1).

**Fig. 2.**
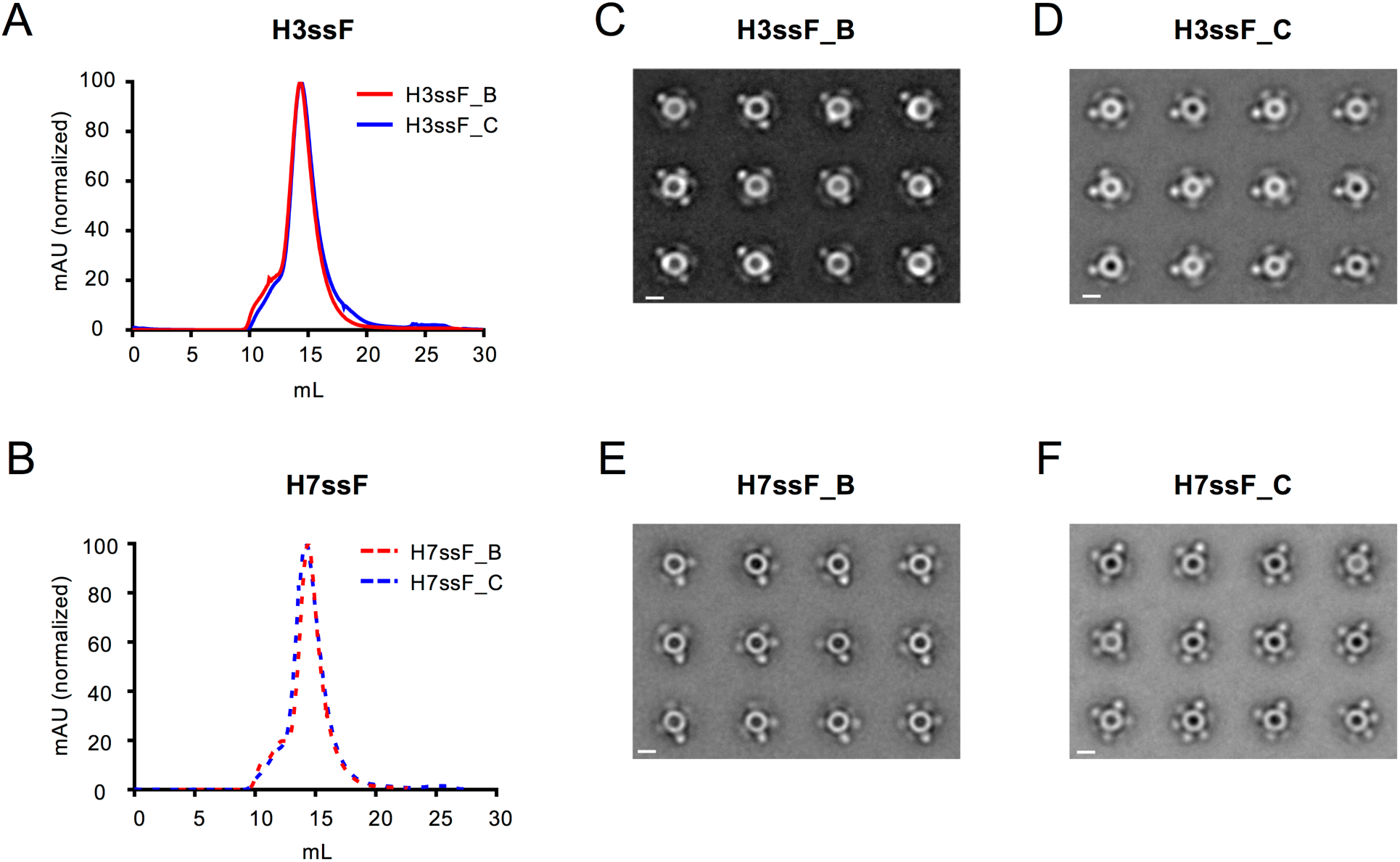
Physical characterization of group 2 HA stem nanoparticles. Gel filtration chromatograms for lectin-purified H3ssF (A) and H7ssF (B) nanoparticles reveal single peaks. Negative stain electron microscopy 2D class averages for H3ssF (C-D) and H7ssF (E-F) B and C variants demonstrate the formation of particles with a visible arrangement of HA stem trimers projecting from hollow spheres. White bars represent 10 nm size markers.

### Thermostability of H3ssF and H7ssF nanoparticles

We next investigated immunogen thermostability using differential scanning calorimetry (DSC). DSC measurements revealed thermal melting temperatures (*T*_*m*_) for the earliest transitions ranging from 56.6 to 66.6°C for H3ssF variants and 56.7 to 61.6°C for H7ssF variants (Table 1 and Fig. S5A-B). Three other higher temperature transitions were consistently observed at approximately 78, 96, and 106°C for all the immunogens tested. In contrast, the ferritin nanoparticle alone had a primary transition at 105.7°C and a minor one at 84°C. Since the earliest transitions were most variable, we hypothesized that these were related the HA stem structures.

The three H3ssF designs with the highest *T*_*m*_s of 66.2-66.6°C had either the seven residue β-hairpin extension disulfide-bonded to helix C (H3ssF_B and H3ssF_C) or the same loop with one glycine removed just after the cysteine (H3ssF_D). Glycine removal just prior to the cysteine (H3ssF_E) was less well tolerated thermodynamically. Curiously, H3ssF_H with the same hairpin loop as H3ssF_B and H3ssF_C, but designed to contain an additional disulfide bond to stabilize the HA1 region of the stem had the lowest *T*_*m*_ of 56.6°C, indicating this variant had less than optimal stability. Design H3ssF_G, engineered to have an internal disulfide bond in the β-hairpin extension had a *T*_*m*_ temperature 6°C lower than H3ssF_B and H3ssF_C. Incorporation of Glu into the two Ala positions in the five-residue helix A extension had little effect on stability (H3ssF_C), but Glu incorporation to replace the last two residues decreased the *T*_*m*_ by 1°C (H3ssF_F). For the five immunogen designs (B-F) made in both H3 and H7 formats, the *T*_*m*_ values were on average 5.9°C higher for the H3ssF relative to H7ssF designs and similar trends were observed between designs. Finally, two additional H7ssF variants not represented in H3ssF, with an Asn incorporated in the HA1 β-hairpin extension (H7ssF_H) and a Pro incorporated into the HA2 GGP loop (H7ssF_I), did not significantly alter the *T*_*m*_. While improved thermostability will facilitate vaccine transport and storage, increasing rigidity and stability of antigens can sometimes be counterproductive for immunogenicity (36). Therefore, multiple designs with varying levels of DSC-determined stability were advanced for further evaluation.

### Antigenicity of H3ssF and H7ssF nanoparticles

Biolayer interferometry (BLI) was used to measure binding kinetics and affinities of nanoparticles to antigen-binding fragments (Fabs) of bNAbs MEDI8852, CT149 and CR8020 (Table 2 and Figs. S6-S7). To avoid avidity effects, the nanoparticles were immobilized to anti-Fc-decorated sensors using the bNAb CR9114 IgG and monovalent Fabs were used as the analyte. CR9114 Fab was observed to recognize H3ssF and H7ssF immunogens with affinities of 94 and 48 nM respectively (Table 2 and Fig. S6E). However, given the bivalent nature of the CR9114 IgG and the 24 epitope sites on each nanoparticle, the avidity of this was complex was more than enough to stably immobilize nanoparticles to the sensor for measurements of high affinity bNAb Fabs.

**Table 2.** Antigenicity of the H3ssF and H7ssF nanoparticles as determined by BLI

| Design | CT149<sup>a</sup> |  |  | MEDI8852<sup>a</sup> |  |  | CR8020<sup>a</sup> |  |  | CR9114<sup>a</sup> |  |  |
| --- | --- | --- | --- | --- | --- | --- | --- | --- | --- | --- | --- | --- |
| | $K_D$ (nM) | $k_{on}$ ($M^{-1}s^{-1}$) | $k_{off}$ ($s^{-1}$) | $K_D$ (nM) | $k_{on}$ ($M^{-1}s^{-1}$) | $k_{off}$ ($s^{-1}$) | $K_D$ (nM) | $k_{on}$ ($M^{-1}s^{-1}$) | $k_{off}$ ($s^{-1}$) | $K_D$ (nM) | $k_{on}$ ($M^{-1}s^{-1}$) | $k_{off}$ ($s^{-1}$) |
| H3ssF_A | ND<sup>b</sup> | ND | ND | ND | ND | ND | ND | ND | ND | ND | ND | ND |
| H3ssF_B | NB<sup>c</sup> | NB | NB | 0.7 | $2.9 \times 10^5$ | $2.2 \times 10^{-4}$ | 20.9 | $2.8 \times 10^5$ | $5.8 \times 10^{-3}$ | ND | ND | ND |
| H7ssF_B | 7.0 | $4.2 \times 10^5$ | $3.0 \times 10^{-3}$ | 1.0 | $2.6 \times 10^5$ | $2.6 \times 10^{-4}$ | NB | NB | NB | ND | ND | ND |
| H3ssF_C | 21.3 | $8.1 \times 10^5$ | $1.7 \times 10^{-2}$ | 0.8 | $3.0 \times 10^5$ | $2.4 \times 10^{-4}$ | 23.3 | $2.4 \times 10^5$ | $5.5 \times 10^{-3}$ | 93.6 | $7.0 \times 10^4$ | $6.5 \times 10^{-3}$ |
| H7ssF_C | 0.9 | $6.0 \times 10^5$ | $5.3 \times 10^{-4}$ | 1.0 | $2.7 \times 10^5$ | $2.6 \times 10^{-4}$ | NB | NB | NB | 47.9 | $1.0 \times 10^5$ | $4.8 \times 10^{-3}$ |
| H3ssF_D | NB | NB | NB | 1.1 | $2.9 \times 10^5$ | $3.2 \times 10^{-4}$ | 21.7 | $2.8 \times 10^5$ | $6.0 \times 10^{-3}$ | ND | ND | ND |
| H7ssF_D | 9.0 | $3.7 \times 10^5$ | $3.3 \times 10^{-3}$ | 0.9 | $2.7 \times 10^5$ | $2.3 \times 10^{-4}$ | NB | NB | NB | ND | ND | ND |
| H3ssF_E | NB | NB | NB | 1.2 | $3.2 \times 10^5$ | $3.7 \times 10^{-4}$ | 18.7 | $3.0 \times 10^5$ | $5.7 \times 10^{-3}$ | ND | ND | ND |
| H7ssF_E | 8.2 | $3.7 \times 10^5$ | $3.0 \times 10^{-3}$ | 0.8 | $2.9 \times 10^5$ | $2.4 \times 10^{-4}$ | NB | NB | NB | ND | ND | ND |
| H3ssF_F | ND | ND | ND | ND | ND | ND | ND | ND | ND | ND | ND | ND |
| H7ssF_F | 2.7 | $6.7 \times 10^5$ | $1.8 \times 10^{-3}$ | 0.9 | $2.9 \times 10^5$ | $2.4 \times 10^{-4}$ | NB | NB | NB | ND | ND | ND |
| H3ssF_G | NB | NB | NB | 0.9 | $3.0 \times 10^5$ | $2.6 \times 10^{-4}$ | 17.9 | $2.7 \times 10^5$ | $4.8 \times 10^{-3}$ | ND | ND | ND |
| H3ssF_H | 22.2 | $8.2 \times 10^5$ | $1.8 \times 10^{-2}$ | ND | ND | ND | ND | ND | ND | ND | ND | ND |
| H7ssF_I | 7.2 | $4.1 \times 10^5$ | $2.9 \times 10^{-3}$ | 1.0 | $2.6 \times 10^5$ | $2.7 \times 10^{-4}$ | NB | NB | NB | ND | ND | ND |
| H7ssF_J | 9.8 | $3.5 \times 10^5$ | $3.4 \times 10^{-3}$ | 1.9 | $2.5 \times 10^5$ | $4.9 \times 10^{-4}$ | NB | NB | NB | ND | ND | ND |
| H3 HA | 42.2 | $7.5 \times 10^4$ | $3.2 \times 10^{-3}$ | 1.2 | $1.3 \times 10^5$ | $1.5 \times 10^{-4}$ | ND | ND | ND | ND | ND | ND |
| H7 HA | 3.7 | $3.6 \times 10^5$ | $3.1 \times 10^{-3}$ | 1.2 | $1.1 \times 10^5$ | $1.3 \times 10^{-4}$ | ND | ND | ND | ND | ND | ND |

<sup>a</sup>Fab
<sup>b</sup>ND, not determined
<sup>c</sup>NB, no detectable binding

The first two antibodies (MEDI8852 and CT149) recognize similar HA stem epitopes centered on the HA2 helix A and fusion peptide loop supersite and represent two multi-donor classes of cross-group bNAbs (HV6-1+HD3-3 class and HV1-18 QxxV class, respectively) recently identified in healthy human adults vaccinated with H5N1 (12, 14, 15). CR8020 is a group 2 specific bNAb that recognizes a less conserved, more membrane proximal HA stem epitope (11). The HV6-1+HD3-3 class monoclonal antibody (mAb) MEDI8852 recognized all H3ssF and H7ssF nanoparticles as well as full length H3 and H7 HA trimers with similar nanomolar affinities and kinetics (Table 2 and Fig. S6A-B). The HV1-18 QxxV class mAb CT149 also recognized all H7ssF nanoparticles with approximately 1-10 nanomolar affinity. Of these, H7ssF_C was recognized with approximately 10-fold greater (0.9 nM) and a 10-fold slower off rate (5.3 × 10^−4^s^−1^) than the other H7ssF nanoparticles. However, among the H3ssF nanoparticles, only H3ssF_C and H3ssF_H were recognized by CT149, with affinities of 21.3-22.2 nM; the other four H3ssF immunogens showed no measurable affinity to CT149 (Table 2 and Fig. S6). This was surprising since each of these designs were initially chosen based on strong binding to CT149 in ELISA assays. Furthermore, ELISAs performed on purified H3 and H7 immunogens for designs B and C also indicated high affinity to each of the group 2-reactive HA stem antibodies tested (Fig. S7A). This apparent disparity between BLI and ELISA was potentially due to utilization of bivalent IgGs in ELISAs as opposed to monovalent Fabs in BLI assays, which provides an avidity advantage for apparent binding affinity of up to two orders of magnitude (37–39). Consistent with this idea, we observed in ELISAs that the Fab form of CT149 did not bind to H3ssF_B or H3ssF_C and bound only modestly to H7ssF_B and H7ssF_C (Fig. S7B). Moreover, ELISAs used a higher density of immobilized antigen than BLI, and there are inherent differences in the methodologies of the two assays. Full length H3 and H7 HA trimers were observed to bind CT149 with affinities and kinetics consistent with the C and H versions of the H3ssF and H7ssF nanoparticles (Fig. S6C-D). Interestingly, the nanoparticles with greatest affinity for CT149, H3ssF_C, H3ssF_H and H7ssF_C, were each designed to have two outwardly facing Glu residues at the C-terminal end of helix A whereas the other designs had Ala at these positions.

The group 2-specific HA stem bNAb CR8020 recognized all of the tested H3ssF nanoparticles with affinities of 17.9-23.3 nM, but did not recognize any of the H7ssF nanoparticles (Table 2 and Fig. S7C). This is consistent with the previously reported lack of CR8020 binding to full length A/Shanghai/2/2013 (H7N9) HA, the same strain used in the H7ssF designs (40). The CR8020 epitope is not completely conserved even within group 2 HAs as there are six epitope mutations between the A/Hong Kong/1/1968 (H3N2) HA and A/Shanghai/2/2013 (H7N9) HA.

Newly recombined, naïve B cell receptors (BCRs) need to first recognize an immunogen to initiate affinity maturation and B cell proliferation (41). Therefore, we also used BLI to assess the relative affinities of three sets of mature stem bNAbs and their inferred UCAs to HA stem nanoparticle immunogens (Fig. S7D). The UCAs were each inferred phylogenetically from multiple clonal family members and include unmutated CDRH3 loops (Fig. S8). Two of these (54-1G07 and 09-1B12) were from the human multi-donor HV6-1+HD3-3 class and HV1-18 QxxV class, respectively (13). The third antibody, 04-1D02 was from an HV3-53+HD3-3 lineage. Measurable H3ssF and H7ssF immunogen recognition by inferred UCAs for each of the three antibodies was not considerably different from the binding of mature forms of the antibodies or the CR9114 positive control (Fig. S7D). This is interesting considering that there are 2-5 unmutated CDRH3 residues in these UCAs (Fig. S8) and crystal structures of mAb complexes with HA show considerable CDRH3 binding for members of the HV1-18 QxxV and HV6-1 HD3-3 multi-donor antibody classes (12, 14, 15).

### Immunogenicity of H3ssF and H7ssF nanoparticles

To evaluate the immunogenicity of the group 2 HA stem nanoparticles, we immunized mice three times intramuscularly at four week intervals with the B and C designs of H3ssF and H7ssF. Both H3ssF and H7ssF elicited robust homosubtypic antibody responses against H3 and H7, respectively (Fig. 3A-B). Similarly, strong homotypic H3ssF and H7ssF neutralizing antibody responses were elicited against H3N2 A/Wisconsin/67/2005 and H7N9 A/Anhui/1/2013 pseudoviruses, respectively (Fig. 3C-D). For both H3 and H7, B and C designs elicited similar levels of antibody responses (Fig. A-D). Following homosubtypic challenges, H3ssF-vaccinated mice were fully protected against H3N2 A/Philippines/2/1982 influenza (shown with H3ssF_C); likewise, H7ssF vaccination fully protected mice from H7N9 A/Anhui/1/2013 (shown with H7ssF_C). Overall, these animal studies demonstrated robust immunogenicity and protective efficacy of our group 2 HA stem nanoparticles against homotypic viruses, with limited cross-recognition of heterosubtypic strains by elicited sera. Although the stabilized group 2 HA stem antigens are immunogenic and protective, murine species tested do not have the appropriate B cell repertoire to evaluate the potential for eliciting bNAbs with specific characteristics similar to known classes of human mAbs with cross-subtype binding (12, 42, 43).

**Fig. 3.**
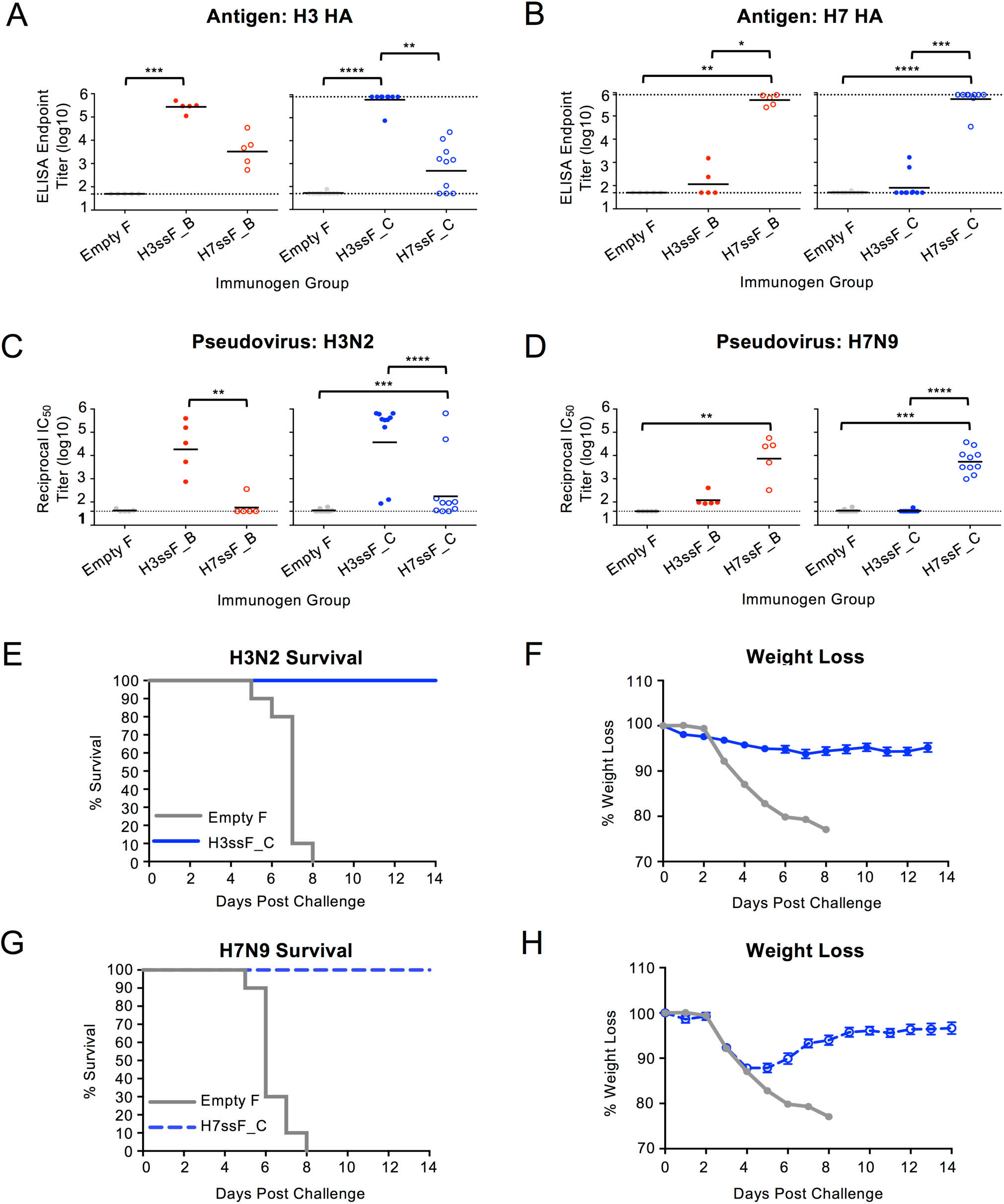
Immunogenicity of group 2 HA stem nanoparticles in mice. BALB/c mice were immunized with H3ssF or H7ssF. B and C design groups were tested in separate experiments. Serum was assessed for binding antibodies to H3 A/Hong Kong/1/1968 HA (A) and H7 A/Anhui/1/2013 HA (B) and neutralizing antibodies against H3N2 A/Wisconsin/67/2005 (C) or H7N9 A/Anhui/1/2013 pseudovirus (D). H3ssF_C and H7ssF_C immunized mice were challenged with lethal doses of H3N2 A/Philippines/2/1982 or H7N9 A/Anhui/1/2013, respectively. Survival (E,G) and weight loss (F,H) were recorded post-challenge. Dotted lines in A-D represent assay limits of detection. One-way ANOVA with Kruskal-Wallis post-test was used to compare mean ELISA and IC_50_ titers between groups. * = p < 0.05, ** = p < 0.01, *** = p < 0.001, **** = p < 0.0001.

### B cell activation by H3ssF and H7ssF nanoparticles

Recent studies have indicated that characterization of *in vitro* engagement of V-gene-reverted BCRs by antigen can be used to assess and rank order vaccine immunogens (43). Therefore, to better understand how the immunogens may be able to engage elements of the human antibody repertoire, Ca^++^ flux assays were used to measure activation of Ramos B cells expressing IgM versions of the inferred UCA of the human antibodies 16.a.26 and 54-1G07 representing two multi-donor classes of cross-group human bNAbs (HV1-18 QxxV class and HV6-1+HD3-3 class, respectively) (12, 13, 43). All eight tested H3ssF and H7ssF designs strongly activated the 54-1G07 UCA BCRs with magnitudes comparable to the anti-IgM control, whereas a group 1 H1ssF negative control resulted in no activation (Fig. 4, left panels). In contrast, only two H3ssF immunogens (H3ssF_C and H3ssF_H) and one H7ssF immunogen (H7ssF_C) were observed to elicit a strong activation of the 16.a.26 UCA BCR (Fig. 4, middle panels). No activation was observed of the HV4-34 group 1-specific 01.a.44 UCA BCR (12) (Fig. 4, right panels). Notably, these results were consistent with the BLI binding data. In both sets of experiments, representatives of the multi-donor HV6-1+HD3-3 class bNAbs engaged all of the tested H3ssF and H7ssF immunogens and the representatives of the multi-donor HV1-18 QxxV class bNAbs engaged only the C and H design immunogens that contain added Glu in the helix A C-terminal extension. Our results suggest that these design C and H designs could potentially activate nascent B cells expressing unmutated versions of multi-donor HV6-1+HD3-3 or HV1-18 QxxV class bNAbs to initiate B cell proliferation and affinity maturation, thereby eliciting cross-group reactive bNAb responses.

**Fig. 4.**
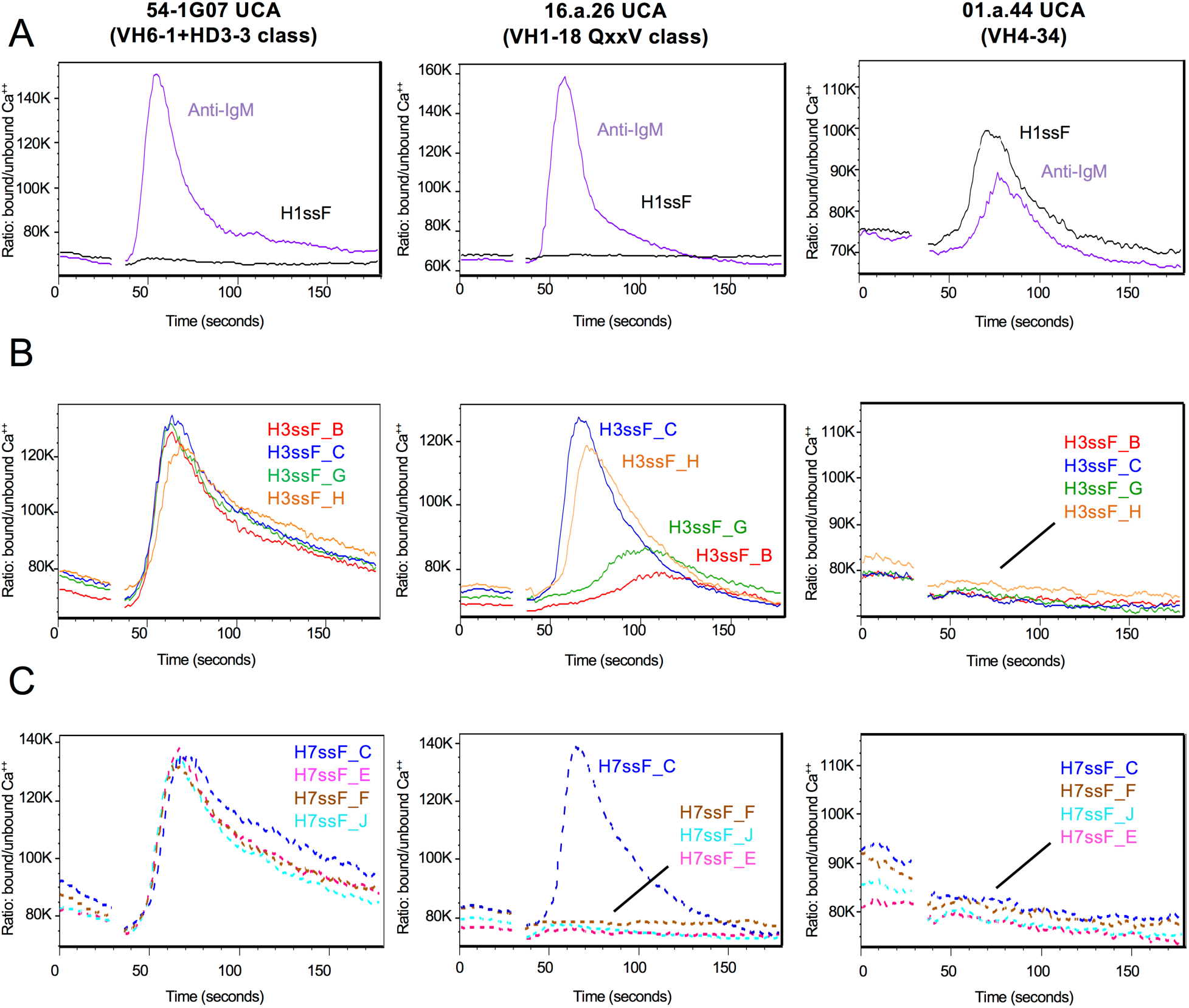
Group 2 HA stem immunogens activate B cells expressing the inferred UCA versions of multi-donor cross-group-reactive human BCRs. The results from Ca^++^ flux assays indicate B cell activation by IgM positive control and a group 1 H1ssF (A), H3ssF variants (B), and H7ssF variants (C). BCR activation is displayed for the group 2-specific 54-1G07 UCA in the left panels, group 2-specific 16.a.26 UCA activation in the middle panels, and the group 1-specific 01.a.44 control BCR in the right panels. Ca^++^ flux was measured by flow cytometry as the ratio (y-axis) of the Ca^++^ bound/unbound states of the Ca^++^ sensitive dye Fura Red over time in seconds (x-axis). Each experiment was repeated three times, and representative curves from one of the repeats are shown for each cell line.

## DISCUSSION

To address the need for influenza immunogens that can elicit broad heterosubtypic responses in humans, particularly against group 2 influenza A, we used an iterative, structure-based design to develop headless group 2 HA stem-only nanoparticle immunogens from both H3 and H7 influenza subtype sequences. Stabilizing the group 2 HA stem was unexpectedly difficult compared to group 1 HA stem antigens. In addition to modifications used to achieve antigenically correct and immunogenic H1ssF candidate vaccines (17), multiple combined protein engineering strategies including helix stabilization, loop optimization, and side chain repacking, were required to achieve the first lead design candidate. Without all three stabilization strategies, the HA stem trimers were not stable enough to be expressed and displayed on self-assembling ferritin nanoparticles. This suggests that screening multiple stabilization methods concurrently may be necessary to design vaccine antigens from metastable proteins or proteins that have had significant portions removed.

Ultimately, H3ssF and H7ssF immunogen designs were identified that provided uniform, thermostable nanoparticles recognized by HA stem-directed bNAbs with subnanomolar to nanomolar affinity. Additionally, H3ssF and H7ssF were immunogenic in mice, eliciting robust homotypic neutralizing antibody responses that fully protected mice against homosubtypic lethal challenges.

The unique antibody repertoire of humans makes it difficult to predict or model the human immune response of candidate influenza vaccines using currently available animal models such as mice (42, 44) or ferrets, particularly given the specific classes of human V-, D- or J-gene combinations associated with broadly cross-reactive immune responses across multiple individuals (12).Therefore, instead of relying on a mouse model to assess potential breadth of antibody response, we utilized *in vitro* Ca^++^ flux assays (43) to evaluate the ability of group 2 HA stem immunogens to activate B cells expressing IgM BCRs comprising inferred UCA forms of the human, cross-group-reactive, HA stem-directed bNAbs from two multi-donor classes (12). Germline targeting with vaccines designed to initiate specific antibody lineages is a new concept being tested with HIV antigens, and not yet proven as a way of achieving broad immunity in humans. Even when transgenic animals are used for preclinical evaluation, the artificial constrains imposed may mitigate correlations with human immune system. Furthermore, since this type of strategy is intended to enhance *de novo* antibody responses it may be particularly difficult to evaluate in pre-immune adults and may require clinical evaluation in naïve children to demonstrate proof-of-concept.

Consistency with the BLI binding results informed the choice to focus on immunogens H3ssF_C, H3ssF_H and H7ssF_C, each of which were designed to have two outwardly facing surface Glu (at positions *i* and *i*+3) at the C-terminus of helix A. The first Glu occurs at a position (HA2 57) that naturally has a Glu in most human H3 strains before 2002 and the majority of human H7 and H10 strains, though it is not in a helical conformation in full length HA. Modeling with CT149 suggests the two Glu residues might interact with light chain CDRL1 and CDRL2 loops (Fig. S9).

The observation that H3ssF and H7ssF variants were able to activate inferred UCA versions of cross-reactive BCRs suggest these immunogens could potentially induce broad cross-group multi-donor antibodies in humans. Due to their broad heterosubtypic breadth (12, 14, 15), such antibodies could potentially protect against pandemic strains in addition to drifted seasonal strains and preclude the need for annual vaccine reformulations. Furthermore, group 2 HA immunogens based on conventional H7N9 monovalent split product formulations have been shown to induce more diverse lineages of cross-group-reactive HA stem antibodies than conventional group 1 H5N1 immunogens in humans (13). Immunological imprinting from earlier or first exposures has also been suggested to play a significant role in shaping the subsequent immune response to influenza (45, 46). Thus, in influenza-naïve individuals, the order of antigen priming may influence the composition of the influenza-specific B cell repertoire. Current vaccine approaches combine multiple strains of influenza in a single formulation. While this may be an effective strategy in adults with pre-existing immunity from prior infections and vaccinations, it is possible that using an influenza A group 2 HA stem-based vaccine as the primary antigen exposure in children could potentially imprint or prime for broader cross-group responses later in life. Moreover, since titers of anti-HA stem antibodies increase with age and are inversely correlated with influenza infection rates, any early robust induction of broad, cross-reactive HA stem antibodies has the potential to mitigate the relative high incidence of severe influenza disease in children in addition to augmenting protection from influenza later in life (47). For adults and elderly who have likely already encountered influenza, the group 2 stem immunogens could selectively boost antibody responses with greater breadth by not engaging otherwise predominant group 1 HA stem-specific HV1-69 antibodies (48, 49). These group 2 immunogens complement our earlier development of group 1 HA stem immunogens which were demonstrated to elicit within-group heterosubtypic protection in mice and ferrets (17). Group 1 HA stem could potentially be used as a boost for the group 2 HA stem immunogens to better focus the antibody response to the neutralization-sensitive epitope centered on helix A. Finally, group 1 and 2 HA stem immunogens could also be given together in a manner analogous to the current multivalent inactivated influenza vaccines (IIV).

Another approach for inducing an HA stem antibody response involves immunizing sequentially with heterologous chimeric HA immunogens comprising two or more HAs with the same stem, but head regions from different subtypes. This strategy has elicited heterosubtypic group 1 and group 2 protection in animal models (50–53). However, the exotic HA heads may still be immunodominant and potentially interfere with or limit HA stem-specific responses as it has been shown that transient vaccine-induced boost of the stem-directed antibody responses can be replaced by immunodominant head-directed responses over time in H5 vaccine trials in humans (49, 54). From a practical clinical perspective, careful records would need be maintained to assure the correct order of administration of heterologous vaccine products.

Although bacterially-produced HA stem-only immunogens from H1, H5, H3 and H7 subtypes have also been reported to protect mice from H3 or H7 influenza, complete protection has only been observed for some homotypic strains (55–57), indicating that the breadth of response is limited. The immunogens in our study more closely mimic the native H3N2 and H7N9 HA stems in that they include native N-linked glycans as well as the membrane proximal region of the HA stem that are both missing in the bacterially-expressed designs. Eukaryotic expression of HA stem immunogens may be of particular importance because each native HA stem trimer has nine glycans and glycans have been observed to guide antibody binding by either masking certain epitopes (58, 59) or participating as part of an epitope surface (60). Indeed, many cross-group-reactive HA stem-directed bNAbs are immediately proximal to the HA2 N38 glycan when recognizing group 2 HA (12). Interestingly, the HA stem-directed bNAbs most frequently elicited from group 1 influenza, which does not have N38 glycans, are of the HV1-69 class (48), that is generally incompatible with the N38 glycan, and therefore not cross-group-reactive. Thus, group 2 HA stem immunogens bearing native glycans may provide a selective advantage for eliciting broad influenza antibody responses. However, to date no experimental data has shown glycosylated hemagglutinin immunogens to be more effective than nonglycosylated versions. A recently reported particulate vaccine comprising nonglycosylated, HA stem crosslinked to desolvated tandem M2e segments has been shown to elicit broad cross-group protection in mice (61). However, these particles comprised proteins derived from multiple host species, strains and subtypes of influenza with the goal of accumulating breadth as opposed to the approach herein of attempting to achieve breadth through targeting conserved antigenic supersites.

In summary, we have designed group 2 headless HA stem trimers with conformational and antigenic fidelity to native HA. These antigens were immunogenic in mice and provided protection against a lethal homosubtypic challenge. Since mice lack critical genetic elements found in human bNAbs targeting the HA stem, we used a B-cell receptor activation assay to demonstrate the potential for eliciting broadly reactive human antibody responses. Based on activation of B cells expressing inferred UCA versions of human antibodies associated with broad cross-neutralizing activity, candidate vaccines with antigens displayed on self-assembling ferritin nanoparticles are now being manufactured for clinical evaluation. Additional animal studies will be performed in parallel, including testing in mice expressing specific human immunoglobulin genes and in nonhuman primates. These immunogens represent a step towards achieving the goal of a vaccine that can elicit protective immunity against drifted seasonal and pandemic strains of influenza A virus.

## MATERIALS AND METHODS

### Structure-based design

The crystal structure of A/Finland/486/2004 (H3N2) HA (PDB IB 2YP2) was used as a template for subtype H3 and the crystal structure of A/Shanghai/2/2013 (H7N9) HA (PDB ID 4LN6) was used as a template for subtype H7. The complete sequences for A/Finland/486/2004 HA and A/Shanghai/2/2013 (H7N9) HA including the signal sequence regions not included in the crystal structures were obtained from GISAID accession EPI397685 and Genbank accession YP_009118475.1 respectively. The change in free energy from point mutations were estimated using the program DDG_MONOMER (26) from the ROSETTA suite. Loops were modeled with LOOPY (62), superpositions were performed using UCSF Chimera (63) or LSQMAN (64), and energy minimization of final models was performed using RELAX (65) from the ROSETTA suite. The graphics programs PyMOL (66) and UCSF Chimera (63) were used for detailed visual inspection of structures and the generation of structural figures.

### Phylogenetic tree generation for HA sequences

Maximum likelihood trees were created using the program MEGA7.0 (ref (67)) from 18 representative HA sequences (13) aligned with Muscle (68). The stem regions of HA sequences were defined as residues HA1 10-42, 314-329 and HA2 1-59 and 93-174 (H3 numbering (21)).

### Antigenic screening of nanoparticle immunogen designs

Most designs were initially screened in a high throughput ELISA format as described previously (23). However, the contributions of individual mutations within multiple-mutation designs were also assessed by expressing transmembrane versions of H3 HA stem containing the native C-terminal HA sequence starting with HA2 175 (SVELKSGYKDWILWISFAISCFLLCVALLGF IMWACQKGNIRCNICI) on the surface of HEK293F cells and measuring recognition of bNAbs FI6v3 and CR8020 using flow cytometry. Since HA stem was more readily expressed in this format with less stabilization required, this strategy enabled a more sensitive evaluation of HA stem mutations than the nanoparticle format. Promising designs that exhibited greater FI6v3 and CR8020 binding were subsequently expressed in the nanoparticle format for final evaluation.

### Expression and purification of immunogens and antibodies

All H3ssF and H7ssF nanoparticles and FI6v3, CT149, CR8020, CR6261, MEDI8852, 315-53-1F12, 54-1G07, 04-1D02 and 09-1B12 mAbs were expressed in 293 Expi cells (Life Technologies) and purified by affinity chromatography (*Galanthus nivalis* lectin for the nanoparticles and protein A for mAbs), using previously described methods (13, 17). Assembly of nanoparticles was assessed by gel filtration with a Superose 6 Increase 10/300 GL column (GE Healthcare). To make MEDI8852 and CR8020 Fabs, IgG was digested with endoproteinase Lys-C (New England Biolabs) overnight at room temperature (RT). To make CT149 Fab, CT149 IgG containing an HRV-3C cleavage site engineered in the heavy chain hinge region was digested using HRV-3C enzyme overnight at RT. The reactions were assessed by SDS-PAGE and upon completion, the reactions were quenched by addition of protease inhibitor cocktail (Millipore-Sigma) and the Fc and Fab were separated by Protein A affinity chromatography.

### ELISA binding assays

ELISA assays were done as previously described (17). Briefly, ELISA plates were coated with 100 ng/well H1ssF, H3ssF, or H7ssF antigens overnight at 4°C. The plates were then incubated with IgG or Fab forms of mAbs with dilutions starting at 10 μg/ml. For Fab, anti-CH1 mAb was then bound at 10 μg/mL. For testing of mouse sera, H3 HA and H7 HA antigens represented A/Hong Kong/1/1968 HA and A/Anhui/1/2013, respectively. Mouse sera serial dilutions started at 1:50. Appropriate HRP-conjugated secondary antibodies were used to detect primary antibody followed by a colorimetric detection assay. Linear regression analysis of the absorbance values (OD 450 nm) was completed using GraphPad Prism software. Serum endpoint titers were determined to be the serum dilution that resulted in four-fold increase in OD value above background. One-way ANOVA with Kruskal-Wallis post-test was used to compare mean endpoint titers between groups. * = p < 0.05, ** = p < 0.01, *** = p < 0.001, **** = p < 0.0001.

### Immunoprecipitation assay

Magnetic protein G Dynobeads™ (Thermo Fisher) were used to capture FI6v3 mAb, per manufacturer’s protocol, and allowed to incubate for 1 hour at RT. FI6v3-bound Dynobeads™ were washed 2 times with 1X PBS and incubated with supernatants from H3ssF or H7ssF expressed in HEK293 cells or purified control protein, for 1 hour at RT. Beads were washed 2 times with 1X PBS and heated to 95 °C for 10 minutes to detach bound protein. Proteins were then analyzed by SDS-PAGE.

### Biolayer interferometry binding assays

All biosensors were hydrated in PBS prior to use. CR9114 IgG (10 µg/ml diluted in 1% BSA-PBS) were immobilized on AHC biosensors through conjugated anti-human Fc antibody (fortéBio). After briefly dipping in assay buffer (1% BSA-PBS), the biosensors were dipped in various H3ssF and H7ssF constructs (10 µg/ml) to capture nanoparticles. The biosensors were then equilibrated in assay buffer for 1 min before dipped in a 2-fold dilution series of CR8020, CT149 or MEDI8852 Fab for 5 min followed by dipping in assay buffer to allow Fab to dissociate from nanoparticles for 5 min. All assay steps were performed at 30°C with agitation set at 1,000 rpm in the Octet HTX instrument (fortéBio). Correction to subtract a baseline drift was carried out by subtracting the measurements recorded for a sensor incubated with no Fab. Data analysis and curve fitting were carried out using Octet analysis software (version 9.0) as previously described (69, 70). Experimental data were fitted with the binding equations describing a 1:1 interaction. Global analyses of the complete data sets assuming binding was reversible (full dissociation) were carried out using nonlinear least-squares fitting allowing a single set of binding parameters to be obtained simultaneously for all concentrations used in each experiment.

### Electron microscopy analysis

Samples were diluted to ∼0.02 mg/ml with buffer containing 10 mM HEPES, pH 7.0, and 150 mM NaCl and adsorbed to freshly glow-discharged carbon-film grids for 15 seconds. The grids were washed with the same buffer and stained with 0.7% uranyl formate. Images were collected at a nominal magnification of 50,000 semi-automatically using SerialEM (71) on an FEI Tecnai T20 microscope operated at 200 kV and equipped with a 2k x 2k Eagle CCD camera. The pixel size was 0.44 nm. Particles were picked automatically using in-house developed software (YT, *unpublished*). Particle alignment and reference-free 2D classification were performed with SPIDER (72) and Relion 1.4 (ref (73)).

### Differential scanning calorimetry (DSC)

Thermal melting points were determined using a MicroCal VP-Capillary differential scanning calorimeter (Malvern Instruments). All protein samples were in the range of 0.2-0.5 mg/ml and extensively dialyzed against phosphate-buffered saline (PBS). Thermal denaturation was probed at a scan rate of 60°C/h from 30 to 120°C. Buffer correction, normalization, and baseline subtraction procedures were applied before the data were analyzed using the VP-Capillary DSC Automated data analysis software.

### Mouse immunizations

Animal experiments were carried out in compliance with all pertinent US National Institutes of Health regulations and policies. The National Institutes of Health, National Institute of Allergy and Infectious Diseases, Vaccine Research Center Animal Care and Use Committee reviewed and approved all animal experiments. Female BALB/cJ mice aged 6–8 weeks (Jackson Laboratory) were immunized with 2 μg H3ssF or H7ssF, adjuvanted with Sigma Adjuvant System, at 0, 4, and 8 weeks. Mice were inoculated with 100 μL intramuscularly, given as 50 μL into each hind leg. Two weeks after the final immunization, sera were collected for measurement of antibody responses. When appropriate, challenges were performed at 4-8 weeks post-immunization. Mice were inoculated intranasally with a 10xLD_50_ dose H3N2 A/Philippines/2/1982, or a 20xLD_50_ dose H7N9 A/Anhui/1/2013. Weight loss was recorded daily for 14 days post-challenge.

### Pseudovirus neutralization assays

Lentivirus-based pseudoviruses displaying influenza HA and NA were produced as previously described (17). Mouse serum was treated with receptor destroying enzyme (Denka Seiken). Serial dilutions of mouse sera (1:40, four-fold, eight dilutions) were mixed with either H3N2 A/Wisconsin/67/2005 or H7N9 A/Anhui/1/2013 pseudovirus (which were previously titered to target 50,000 RLU), for 30 minutes at RT. The virus/pseudovirus mixture was then added to previously-plated 293A cells (ATCC), in triplicate. After 2 hours, at 37 °C and 5% CO_2_, fresh DMEM supplemented with 10% FBS, 2 mM glutamine, and 1% penicillin/streptomycin was added. Cells were lysed at 72 hours, and luciferase substrate (Promega) was added. Luciferase activity was measured as relative luciferase units (RLU) at 570 nm on a SpectramaxL (Molecular Devices). Sigmoidal curves, taking averages of triplicates at each dilution, were generated from RLU readings; 50% neutralization (IC_50_) titers were calculated considering uninfected cells as 100% neutralization and cells transduced with only virus as 0% neutralization. One-way ANOVA with Kruskal-Wallis post-test was used to compare mean IC_50_ titers between groups. * = p < 0.05, ** = p < 0.01, *** = p < 0.001, **** = p < 0.0001.

### Determination of inferred UCAs for HA stem antibodies

Paired heavy and light chain DNA sequences for 58 members of the 54-1G07 clonal family (315-54-1G07) were isolated from single IgG^+^ B cells, along with 6 additional heavy chain DNA sequences from cells for which light chains could not be amplified (13). A maximum likelihood phylogenetic tree was constructed from concatenated heavy and light chain sequences and the most recent common ancestor (MRCA) of the lineage was inferred using DNAML in SONAR (74). This MRCA had 5 residual nucleotide mutations each in heavy and light chain (resulting in 2 and 5 amino acid changes, respectively) compared to the assigned germline VH and VK genes. To construct the final UCA, these changes were reverted to the presumed germline sequence. The same procedure was used for 57 paired heavy and light chain DNA sequences and 8 unpaired heavy chain DNA sequences from the 09-1B12 clonal family. This MRCA had 1 residual nucleotide mutation (1 amino acid change) in the heavy chain and no residual mutations in the light chain. For the 04-1D02 clonal family, 29 paired heavy and light chain DNA sequences and 1 unpaired heavy chain DNA sequence were used. The inferred MRCA contained 4 residual nucleotide mutations in the heavy chain and 6 nucleotide mutations in the light chains (4 and 5 amino acid changes, respectively). For this clonal family, only changes in VH genes were reverted; the lambda chain of 04-1D02-MRCA was used with the residual mutations included.

For the 16.a.26 clonal family, 93 heavy chain, and 83 light chain sequences and for the 01.a.44 clonal families, 51 heavy chain and 49 light chain sequences were identified from single sorted IgG^+^ B cells. For the heavy chain, a maximum likelihood phylogenetic tree based upon V-gene sequence alone was constructed using Geneious (75), with each tree rooted to the respective germline reference sequence as listed in the IMGT database. The VDJ junction of clonal members with the lowest mutation load was inferred using IMGT V-quest, allowing identification of residues within the V and D and J gene segments most likely subject to somatic mutation. To construct the heavy chain UCA, these changes were reverted to the presumed germline sequences. Due to the lack of precision in estimating N-nucleotide addition, mutations falling at V/D or D/J junctions were not reverted. For the light chains, an analogous process was used for V and J gene segments.

### Generation of B cell lines

B cell lines expressing inferred UCA versions of monoclonal antibodies were generated by transduction of surface IgM-negative Ramos cells as previously described (43). Briefly, DNA encoding for VDJ (heavy chain) and VJ (light chain) immunoglobulin regions was synthesized by GenScript and cloned, along with the consensus human IgM C region, into the pLVX-ZsGreen and pLVX-mCherry expression vectors (both from Clontech) respectively. Each of these vectors was then co-transfected with the lentivirus packaging plasmid psPAX2 and with the VSV-G envelope expressing plasmid pMD2.G (both from Addgene) into HEK293 T cells for formation of lentiviral particles using Lipofectamine 2000 (Thermofisher). Supernatant was harvested 3 days after transfection and cleared by centrifugation before IgM-negative Ramos cells were co-transduced for both heavy and light chains. After 5 days, transduced Ramos cells were enriched by cell sorting for expression of ZsGreen and mCherry using a FACS AriaII interfaced to the FacsDiva software (BD Biosciences). This procedure was repeated until a pure double positive population could be selected. Cell lines were further enriched for high IgM surface expression by using a fluorescently labeled anti-human IgM monoclonal antibody.

### B cell activation assays

The ability of the different particles to induce calcium flux upon BCR engagement was measured in vitro using inferred UCA monoclonal antibodies expressed as surface IgM in Ramos cells as previously described (76). Briefly, 1 million cells (per test) were stained with 0.35 μL Fura Red (Thermo Fisher Scientific) in 100 μL serum-free medium at ambient temperature in the dark for 30 min. After washing, cells were resuspended in 300 μL serum-free medium and heated to 37°C for 3-5 min in a heated water bath before they were acquired on a FACS Symphony interfaced to the FacsDiva software (BD Biosciences). Cells were first acquired for 30 s in the absence of antigen to record baseline intracellular calcium levels and then tubes were removed (leaving the acquisition in progress) to add the antigen and quickly vortexed before placed back on acquisition for a total time of 180 seconds. Nanoparticles were used at the final concentration of 50 nM. An anti-human unlabeled Mouse F(ab’)2 Anti IgM (Southern Biotech) was used (1.5 μg/test) to determine the maximal induction of calcium flux measured by the change between the emission of Fura Red bound to calcium and the one unbound (ratio) over time. The inferred UCA for 01.a.44 was chosen as a negative control since it is observed to bind only to group 1 HAs and not group 2 (not shown). Likewise, the inferred UCAs for 54-1D07 and 16.a.26 are observed to recognize group 2 HAs only (not shown).

### Accession numbers for inferred antibody UCA sequences

The sequences for the inferred UCA versions of the following HA stem antibodies have been deposited into GenBank: 54-1G07 (MH631450 and MH631451), 04-1D02 (MH997406 and MH997407), 09-1B12 (MH997404 and MH997405), 01.a.44 (MK291363 and MK291365) and 16.a.26 (MK291364and MK291366).

## ACKNOWLEDGEMENTS

This work was supported by the Intramural Research Program of the Vaccine Research Center and the Division of Intramural Research, National Institute of Allergy and Infectious Diseases, National Institutes of Health. EM data collection and analyses were funded by federal funds from the Frederick National Laboratory for Cancer Research, National Institutes of Health, under contract HHSN261200800001E, and Leidos Biomedical Research, Inc. (Y.T.). K.S.C.’s research fellowship was partially funded by the Undergraduate Scholarship Program, Office of Intramural Training and Education, Office of the Director, National Institutes of Health. J.C.B., K.S.C., S.M.M., Y.M.H., M.K., L.W., J.R.M. and B.S.G. are named inventors on pending patent applications involving influenza vaccines.

## SUPPLEMENTARY FIGURE LEGENDS

**Fig. S1.** Design of group 1 HA stem nanoparticle immunogens. Ribbon diagrams depict the design of HA stem-ferritin nanoparticles by 1) removing the head region of HA and replacing with glycine-rich linkers, 2) two internal packing mutations and 3) genetically fusing to a ferritin nanoparticle by a three-residue linker. Structures are shown by ribbon diagrams. One monomer of each HA and HA stem trimer is colored blue and mutated regions or residues are colored red. The nanoparticle core is brown.

**Fig. S2**. Sequence alignments of select H3ssF (A) and H7ssF (B) variants with the full length HA ectodomains for A/Finland/486/2004 (H3N2) (H3_FI04) and A/Shanghai/2/2013 (H7_SH13) respectively. Mutations are in red. Residues identical to the full length sequence are represented by dots and deletions represented by dashes. H3N2 numbering is shown above each alignment. For clarity, the HA head regions and the ferritin sequence is not shown. In each construct, the C-terminal HA2 174 residue is connected to ferritin residue Asp5 by a short SGG linker (not shown).

**Fig. S3.** Characterization of H3ssF_A. (A) Superose 6 gel filtration chromatograms for and H3ssF_A (magenta) and H3ssF_B (red) nanoparticles. (B) SDS PAGE analysis of the results from an FI6v3 immunoprecipitation of the supernatants from H3ssF_A expressed in HEK293 cells. Molecular weight standards (MW) are designated as kDa. The band indicative of H3ssF_A is boxed in magenta. H3ssF_null is previous design iteration that did not express. H1ssF is shown as a positive control. (C) Negative stain electron microscopy 2D class averages of gel filtration purified H3ssF_A. The white bar represents a 10nm size marker.

**Fig. S4.** Physical characterization of H3ssF (A-E) and H7ssF (F-J) nanoparticles. Superose 6 gel filtration chromatograms (left panels) for lectin-purified nanoparticles reveal single peaks. Negative stains electron microscopy 2D class averages (right panels) demonstrate the formation of particles with a visible arrangement of HA stem trimers projecting from hollow spheres. White bars represent 10nm size markers for right panels.

**Fig. S5.** Differential scanning calorimetry (DSC) plots for group 2 HA stem immunogens. (A) H3ssF. (B) H7ssF. (C) Ferritin alone. Plots of heat capacity (Cp) versus temperature depicts melting transitions for each protein. The Cp values on the Y-axis are shown with an arbitrary scale.

**Fig. S6.** BLI binding curves for MEDI8852 (A and B) and CT149 (C and D) Fab recognition of H3ssF and H7ssF immunogens respectively. Binding constants and kinetic parameters for each plot are shown in Table 2. (E) Binding of H3ssF_C and H7ssF_C to CR9114 Fab. Nanoparticles were immobilized to the sensor tip by binding to CR9114 IgG coupled by human anti-Fc antibody and HA trimers were immobilized on HIS1K sensors through C-terminal His tags. Data curves are in red; fitting for a 1 to 1 interaction are in black.

**Fig. S7.** Antigenic recognition of group 2 HA stem nanoparticles. (A) ELISA binding of H3ssF and H7ssF designs B and C by six broadly neutralizing HA stem antibodies. (B) Relative recognition of H3ssF (top) and H7ssF (bottom) by IgG and Fab forms of CT149 as measured by ELISA. In both (A) and (B), the nanoparticles were immobilized on the plate. (C) BLI binding curves for CR8020 Fab recognition of H3ssF (left) and H7ssF (right) immunogens. Binding constants and kinetic parameters for each plot are shown in Table 2. (D) Relative antigenic recognition of H3ssF and H7ssF by mature and inferred UCAforms of HA stem human antibodies as measured by BLI. CR9114 and the HA head antibody CH65 served as positive and negative controls respectively. Antibody recognition was determined by Octet and binding relative to CR9114 was plotted on the y axis as a percentage. In (C) and (D), all nanoparticles were immobilized to the sensor tip by binding to CR9114 IgG coupled by human anti-Fc antibody.

**Fig. S8.** Sequence alignments of mature and inferred UCA for mAbs 09-1B12, 16.a.26, 54-1G07, 04-1D02 and 01.a.44. Dots indicate identical residues. CDR loops are boxed according to Kabat numbering and convention. Shading indicates the antibody classes labeled in the upper left of each box.

**Fig. S9.** Model of CT149 interaction with helix A of H3ssF_C. The left panel shows a ribbon diagram model of CT149 (green and brown) bound to H3ssF_C (blue and gray). Helix A from one protomer of the H3ssF_C model was superimposed onto HA helix A (chain B) of the CT149/H3N2 HA crystal structure (PDB entry 4UBD) using PyMOL. For clarity, the H3N2 HA is not shown and only one CT149 Fab (chains C and D) is depicted. The middle panel shows a close-up of the light chain interaction with HA helix A and the right panel shows the same close-up view rotated 90° along the x-axis. For clarity, the middle and right panels show only helix A from one chain of the H3ssF_C model. The two Glu side chains at the C-terminal end of H3ssF_C helix A are shown as stick models.

